# Melanoma-intrinsic NR2F6 activity regulates anti-tumor immunity

**DOI:** 10.1101/2022.09.14.508028

**Authors:** Hyungsoo Kim, Yongmei Feng, Rabi Murad, Victoria Klepsch, Gottfried Baier, Ze’ev A Ronai

## Abstract

Nuclear receptors (NRs) have been implicated in tumor and immune cell regulation, most notably in hormone-dependent cancers. Here we identify a tumor-intrinsic function of the orphan nuclear receptor NR2F6 in regulating anti-tumor immunity. We selected NR2F6 among 48 candidate NRs based on an expression pattern in melanoma patient specimens coinciding with the IFNγ signature, which is associated with positive responses to immunotherapy and favorable patient outcomes. Further delay of melanoma development was achieved upon combination of NR2F6 inhibition with anti-PD1 therapy. NR2F6 loss in B16F10 and YUMM1.7 melanoma cells attenuated tumor development in immune-competent but not -incompetent mice via increased CD8^+^ T cell infiltration and activation. Inhibition of NACC1 and FKBP10, identified here as NR2F6 effectors, phenocopied NR2F6 loss. Remarkably, inoculation of NR2F6 KD melanoma cells into mice genetically deficient in NR2F6 further enhanced tumor growth inhibition. Identifying a tumor-intrinsic function of NR2F6 which complements its tumor-extrinsic role could be thus exploited to develop novel anti-cancer therapies.

## Introduction

The importance of the tumor microenvironment (TME) for cancer development has gained significant traction as we gain insight into cross-talk between TME components, including immune cells, cancer-associated fibroblasts, endothelial cells (i.e., stroma), and tumors (Belli et al., 2018; Friedl and Alexander, 2011; Hanahan and Weinberg, 2011; Kolch et al., 2015; Quail and Joyce, 2013; Spranger and Gajewski, 2018). These interactions rely in part on cues derived from tumor cells (tumor-intrinsic), although changes in TME components (tumor-extrinsic) are also regulated by cytokines and chemokines secreted by tumors or TME components (Labani-Motlagh et al., 2020; Li et al., 2021b; Santos et al., 2021). These factors govern the recruitment and infiltration of a tumor by stromal components and thus define tumor fate. Our understanding of tumor/TME cross-talk has driven the development of novel therapeutic modalities, including the evolution of immune checkpoint therapies (ICT). ICTs target a natural gatekeeping brake, which evolved to prevent self-attack and maintain immune homeostasis and has revolutionized cancer treatment. Blocking an immune checkpoint via ICT often revitalizes the immune system’s ability to eradicate tumor cells (Carlino et al., 2021; Patel and Minn, 2018; Sharma and Allison, 2015a; b). Indeed, ICT is now the first-line therapy for several cancers, as reflected in the growing number of FDA-approved ICT drugs capable of producing durable tumor remission (Vaddepally et al., 2020). Despite these advances, success is still limited, as a sizable percentage of patients are either non-responsive or develop resistance to ICT (Jenkins et al., 2018; Kalbasi and Ribas, 2020; Schoenfeld and Hellmann, 2020; Spranger and Gajewski, 2016; 2018).

Given these limitations, a growing effort is being devoted to understanding mechanisms governing ICT responsiveness as a means to identify factors that could serve as biomarkers to predict a positive ICT response (Anagnostou et al., 2017; Burr et al., 2019; Emran et al., 2019; Jenkins *et al*., 2018; Kim et al., 2020; Li et al., 2021a; Phan and Croucher, 2020). Thus far, responsiveness to ICT is primarily determined by factors such as a tumor’s mutation burden, the ability of T cells to infiltrate a tumor, and tumor responsiveness to IFNγ (Ayers et al., 2017; Grasso et al., 2020; Rizvi et al., 2015; Tumeh et al., 2014). Thus, identifying regulatory components that could be targeted to improve ICT effectiveness and durability remains an important goal and an unmet clinical need.

Factors that define tumor responsiveness to therapy, including ICT, often influence TME activity. Tumor-intrinsic activities can impact the extracellular matrix, stromal and immune components, or interactions among TME components (Labani-Motlagh *et al*., 2020; Li *et al*., 2021b; Santos *et al*., 2021). In addition to the chemokines and cytokines noted above, nuclear hormone receptors (NRs) also modulate tumor cell communication with the TME and may underlie a tumor’s response to external stimuli (including therapy), the microenvironment (nutrient and oxygen availability), and cross-talk with stromal (immune) cells (Dhiman et al., 2018; Font-Diaz et al., 2021; Zhao et al., 2020; Zhao et al., 2019). Here we focus on the latter.

In normal cells, NRs are known to control regulatory pathways that govern proliferation, metabolism, specialized cell functions, and immune cell activities. Likewise, NRs are often deregulated or dysfunctional in pathophysiological conditions, including cancers. Accordingly, NRs are implicated in hormone-dependent cancers (such as breast and prostate cancers) and often serve as markers for patient stratification or as treatment targets (Dhiman *et al*., 2018; Zhao *et al*., 2019).

Recent studies also indicate an emerging role for NRs in anti-tumor immunity. Activation of LXR, for example, induces APOE expression within the melanoma niche to attenuate innate myeloid derived suppressor cells (MDSCs), resulting in more effective inhibition of metastasis (Pencheva et al., 2014; Tavazoie et al., 2018). Conversely, tumor-derived retinoic acid (RA) reportedly induces monocyte differentiation to immune-suppressive tumor-associated macrophages (TAMs), which dampen ICT impact (Devalaraja et al., 2020). Thus, the possibility of controlling NR activity via regulatory ligands or agonists/antagonists has prompted continued efforts to identify NRs that modulate the TME and could be exploited in more effective immunotherapy strategies.

The orphan NR NR2F6, also known as Ear-2 or COUP-TFIII, is a member of the NR2F subfamily structurally related to NR2F1 and NR2F2 proteins (Hermann-Kleiter et al., 2008). In immune cells, NR2F6 has been suggested to be an immune checkpoint candidate based on analysis of its capacity to fine-tune adaptive immunity and repress transcription of the cytokines IL-2, IFNγ, IL-17, and IL-21 (Hermann-Kleiter *et al*., 2008; Hermann-Kleiter et al., 2012; Jakic et al., 2021; Olson et al., 2019). Accordingly, a genetic mouse model with global ablation of NR2F6 exhibits accelerated inflammation, autoimmune phenotypes, and attenuated tumor growth based on increasing augmenting intra-tumoral availability of effector T cells (Hermann-Kleiter *et al*., 2008; Hermann-Kleiter et al., 2015; Jakic *et al*., 2021; Klepsch et al., 2018; Klepsch et al., 2020). Despite these findings, tumor intrinsic NR2F6 expression is less studied. Interestingly, high NR2F6 expression in tumors is linked to pro-tumorigenic phenotypes, including proliferation, invasion, poor prognosis, and resistance to therapy, in colon, ovarian, lung, liver, and breast cancers (Jin et al., 2019; Li et al., 2019; Li et al., 2011; Liu et al., 2017; Wang et al., 2019; Zhang et al., 2020). Moreover, tumor intrinsic NR2F6 function in defining the anti-tumor immune response remains largely unexplored.

Here, we show that tumor intrinsic NR2F6 expression in melanoma cells may mediate immune system evasion. We report that NR2F6 controls expression of genes functioning to suppress anti-tumor immunity. Correspondingly, higher NR2F6 expression in melanoma patient specimens was associated with a less favorable prognosis and poor response to ICT. These findings impact ICT strategies and could provide a basis to stratify patients for therapy based on NR2F6 expression and a justification for systemic targeting of NR2F6 as therapy.

## Results

### Identification of NRs that govern anti-tumor immune responses

To identify NRs that control responses to ICT, we searched for NRs expressed in melanoma tumor cells that may regulate anti-tumor immunity. To do so, we compared transcriptome data from bulk and single-cell RNAseq analysis of patients’ specimens with patient clinical outcomes to identify NRs whose expression correlated with an immune-responsive gene signature. We then correlated their expression with patient survival and favorable responses to ICT (Figure 1A). IFNγ signaling is a well-established immune responsive gene signature, one enriched in melanoma patients that respond to immune checkpoint blockade (Ayers *et al*., 2017): patients whose specimens show high expression of IFNγ signature genes exhibit better response to ICT and overall survival (Ayers *et al*., 2017). We found that of the 48 NRs expressed in human tissues examined, 20 were either positively or negatively correlated with IFNγ-signature genes (Figure 1B). To narrow the candidate list, we asked which of the 20 exhibited a significant correlation with patient survival and responses to ICT and found that 11 showed significant positive or negative correlations with patient outcomes (Figure 1C, upper bars). Of those, 8 were significantly correlated with an ICT response (Figure 1C, lower bars), 5 (NR1H3, NR3C1, RORC, ESR1, ESRRB) exhibited a positive correlation and 3 (NR2F6, NR6A1, ESRRA) a negative correlation with expression of IFNγ-signature genes, prolonged patient survival and a response to immune therapy (Figure 1C). Next, given that NR expression in tumor cells may modulate either immune evasion or anti-tumor immunity, we assessed NR expression in primary melanoma tumor cells using single-cell transcriptomic data (Jerby-Arnon et al., 2018). Among NRs expressed in >10% of melanoma cells were NR1H3, NR3C1, NR2F6, ESRRA (Figure 1D), which are highly expressed in malignant cells (13-51%) and either positively or negatively correlated with the IFNγ-signature (Figure 1E), responses to ICT (Figure 1F) and overall patient survival (Figure 1G). Of these NRs, three (ESRRA, NR1H3, and NR3C1) were also expressed in multiple immune cell types in the TME, while only one, NR2F6, was expressed primarily in melanoma and non-immune stromal cells, including endothelial cells and cancer-associated fibroblasts (Figure S1B-C). We then assessed the function of these four NRs in regulating anti-tumor immunity.

**Figure 1.**
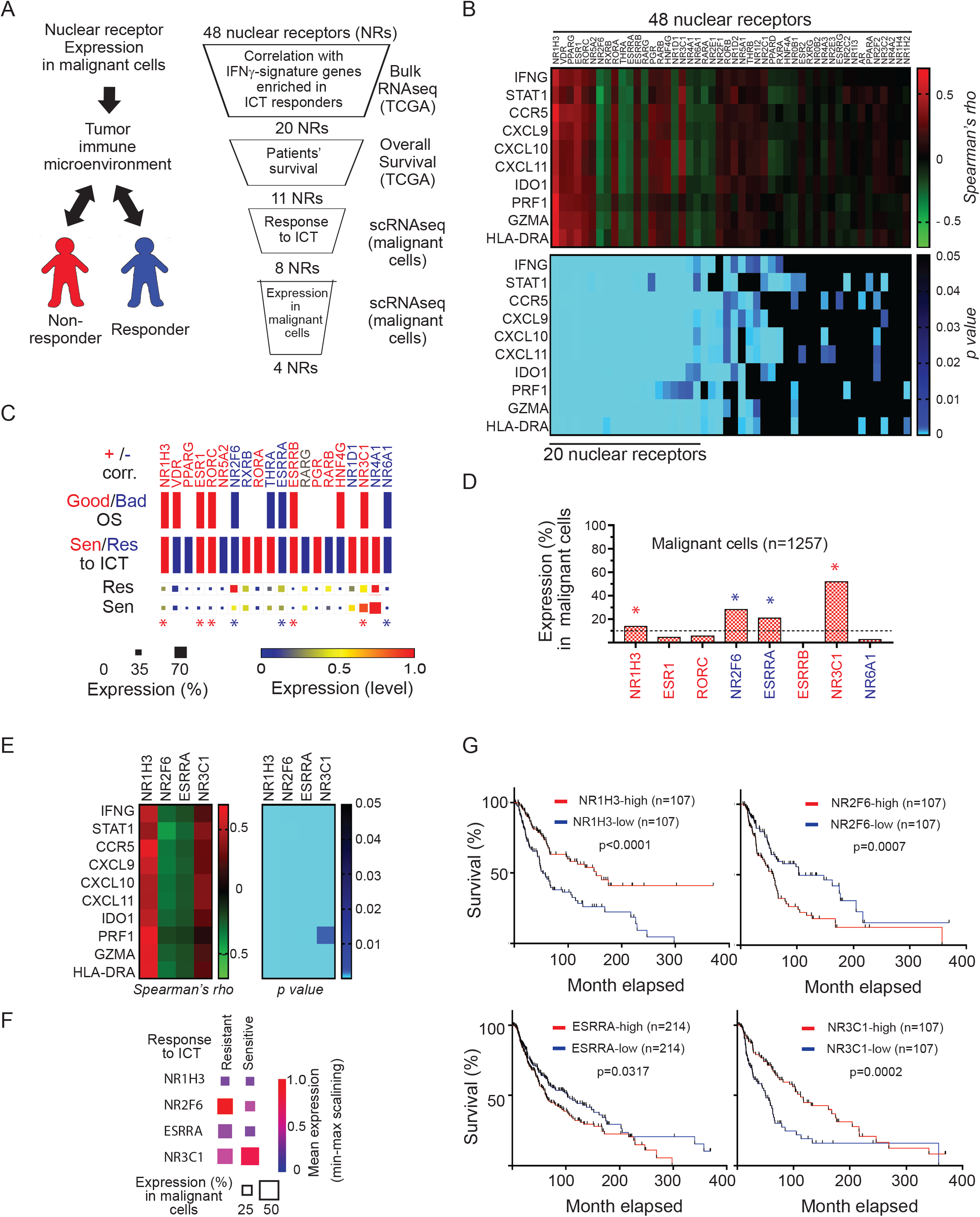
Identification of NRs that may be linked with anti-tumor immunity in melanoma patients. (A) Outline of workflow in the selection process. (B) Heatmaps show Spearman correlation coefficient (upper) and corresponding p values (lower) in comparisons of expression of 48 NRs (X-axis) with 10 IFNγ−signature genes (Y-axis). Twenty NRs showing a significant correlation wi*th* the IFNγ−signature were selected. (C) Those 20 NRs positively (red) or negatively (blue) correlated (Corr.) with IFNγ−signature genes were assessed for potential correlation with patient overall survival (upper panel) and patient responses to ICT, (lower panel). Relevant to overall survival, red and blue bars represent favorable and unfavorable overall survival relative to the expression of corresponding NRs. Red and blue bars relevant to ICT response represent “sensitivity” and “resistance” of patients with high NR-expressing tumor cells to ICT. Squares at the bottom show percent expression and expression level of NRs in tumor cells from patients resistant (upper) or sensitive (lower) to ICT. Eight NRs with a significant correlation with patient response to ICT were selected. (D) Tumor intrinsic expression of those 8 NRs was assessed using scRNAseq data from melanoma patients (Tirosh *et al*., 2016). Four NRs expressed in >10% of tumor cells were either positively or negatively correlated with IFNγ−signature genes (E), response to ICT (F), or overall patient survival (G).

### NR2F6 control of murine melanoma growth requires an intact immune system

We next asked whether altering expression of the four candidate NRs in melanoma cells inoculated in mice would change melanoma responses to ICT using anti-PD1 antibodies (RMP-14 clone) as checkpoint blockers. To this end, we engineered B16F10 cells, which do not respond to ICT (Kim *et al*., 2020), to either knock down (via shRNA) NR2F6 or ESRRA or overexpress NR1H3 or NR3C1. The engineered melanoma cells were then inoculated in mice to follow up their growth. Tumor growth following injection of B16F10 cells overexpressing NR1H3 or NR3C1 (Figure S2A) or deficient in NR2F6 or ESRRA (Figures 2A, S2D) was monitored in immunocompetent C57/BL6 mice to determine whether manipulations activated an anti-tumor immune response and attenuated tumor growth. To exclude possible changes in rate of tumor cell growth following NR manipulation, we first assessed growth of each of the four engineered B16F10 lines in culture. We observed no changes in growth rate in cells overexpressing NR1H3 or NR3C1 or deficient in NR2F6 (Figures 2B, S2B). By contrast, ESRRA knockdown limited growth of B16F10 cells in culture, consistent with observations made in human melanoma cells depleted of ESRRA [DeMap (https://depmap.org/portal/)]. Next, each of these four melanomas engineered cultures was inoculated in mice and tumor formation was monitored with and without ICT. Among the three NRs tested, NR2F6 knockdown in tumor cells notably delayed melanoma growth in mice treated with anti-PD-1 antibodies (Figure 2C), while overexpression of either NR1H3 or NR3C1 had no effect on ICT responsiveness (Figure S2C). These observations suggest that NR2F6 may impact tumor-intrinsic mechanisms that convert B16F10 cells from an immunologically cold to warm status and allow a response to ICT. To directly assess this possibility, we compared growth of B16F10 cells deficient in NR2F6 in immune incompetent versus competent mice. Notably, compared with B16F10 cells with wild-type NR2F6 expression, NR2F6 loss via either shRNA or CRISPR led to 20-60% inhibition of tumor growth in immune-competent (C57/BL6) but not - incompetent (NSG) mice (Figures 2D-H, S2D-E). Accordingly, loss of NR2F6 expression in melanoma YUMM1.7 cells (*Braf*^*V600E*^*/Pten*^*-/-*^*/CDKN2A*^*-/-*^) also attenuated tumor formation in immune-competent but not -incompetent mice (Figure 2I-L). These findings suggest that anti-tumor phenotypes seen following loss of NR2F6 expression require an intact host immune system. Given that tumor-intrinsic NR2F6 loss attenuates B16F10 cell growth, we asked whether ectopic NR2F6 expression would enhance tumor development in mice. Relative to control B16F10 cells, B16F10 cells overexpressing WT or DNA binding-defective mutant of NR2F6 C112S (Liu et al., 2003) were inoculated in mice following their growth. While elevated expression of NR2F6 did not alter tumor growth, expression of the DNA binding-defective mutant, reminiscent to the NR2F6 knock-down, attenuated the degree of tumor growth in mice and extended overall animal survival (Figures S2F-I), suggesting that NR2F6 transcriptional activity is required for tumor-intrinsic changes in anti-tumor immunity. Of note, as in B16F10 cells, ectopic NR2F6 expression in YUMMER1.7 melanoma cells also had no impact on tumor development. However, unlike B16F10 cells, YUMMER1.7 cells are immune-responsive; thus, these observations exclude the possibility that lack of effects on B16F10 melanoma growth after ectopic NR2F6 expression is attributable to insensitivity of these melanoma cells to ICT. We also cannot exclude the possibility that NR2F6 effects require a co-activator, which must be co-expressed.

**Figure 2.**
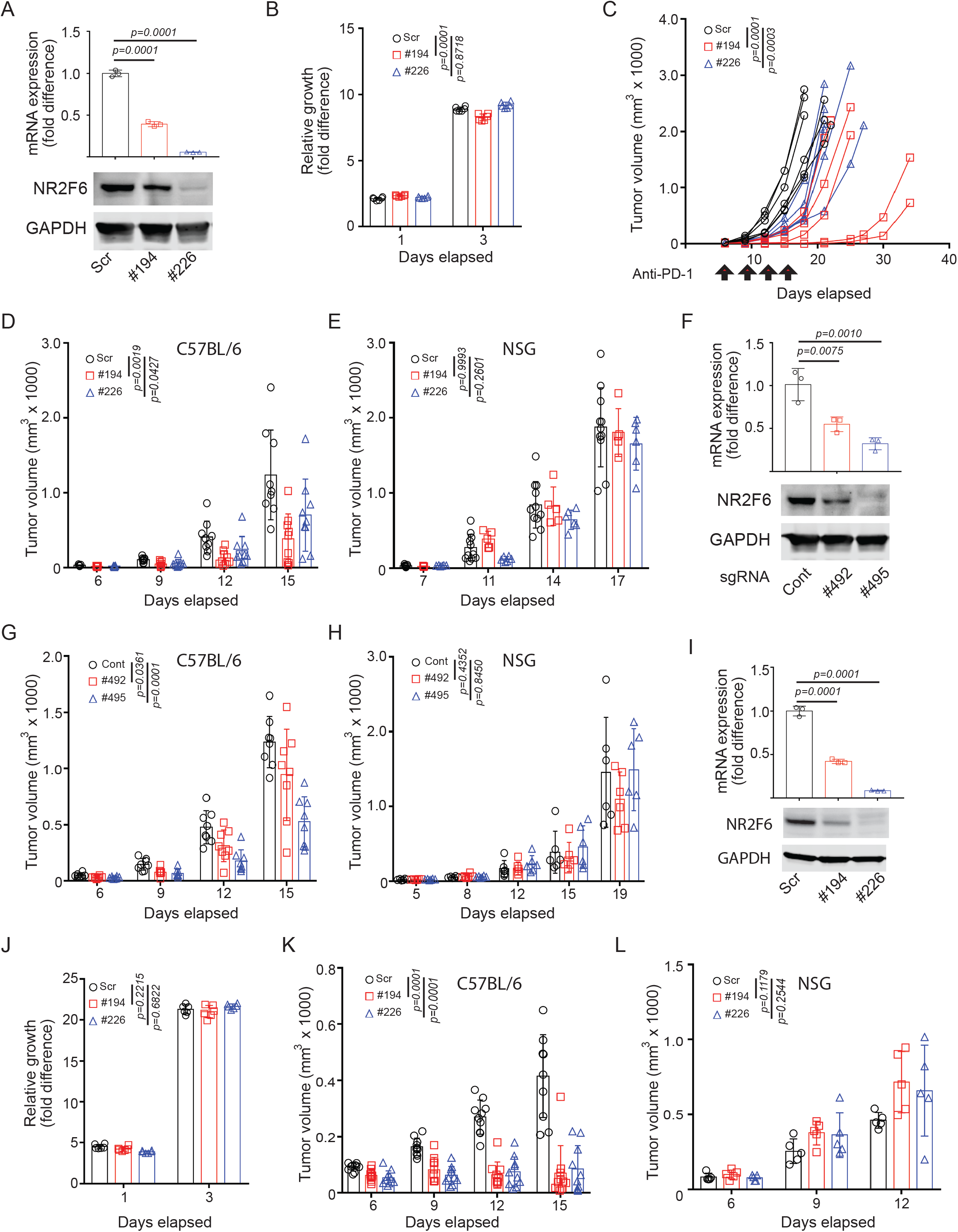
NR2F6 control of murine melanoma growth requires intact immune system. (A) B16F10 cells were transduced with scramble (Scr) shRNA or two shRNAs (#194 and #226) targeting murine Nr2f6. NR2F6 mRNA and protein expression were assessed by qPCR and immunoblotting. (B) Growth of cultured cells described in (A) was assessed *in vitro* using CellTiter-Glo. Relative fold-differences in luminescence on days 1 and 3 were calculated relative to luminescence on day 0, defined arbitrarily as 1. (C) Cells were then inoculated in C57BL/6 mice, which were treated with anti-PD-1 antibody (RMP1-14) at days 6, 9, 12, and 15 (arrows). Tumor volumes were monitored at indicated time points. (D-E) Cells established in (A) were engrafted into C57BL6 (D) or NSG (E) mice, and tumor volumes monitored at indicated time points. (F) Nr2f6 KO B16F10 cells were established by CRISPR using specific sgRNA (#492 and #495). NR2F6 mRNA and protein expression were assessed by qPCR and immunoblotting. (G-H) Cells established in (F) were inoculated in C57BL6 (G) or NSG (Ayers *et al*.) mice, and tumor volumes monitored at indicated time points. (I) YUMM1.7 cells were transduced with scrambled (Scr) shRNA or two shRNAs against Nr2f6 as in (A). mRNA and protein expression were assessed by qPCR and immunoblotting. (J) The growth of cultured YUMM1.7 described in (I) was assessed *in vitro* using CellTiter-Glo. (K-L) Cells established in (I) were then inoculated in C57BL/6 (K) or NSG (L) mice, and tumor volumes monitored at indicated time points. Data are presented as means ± SD. Statistical significance was assessed by two-way ANOVA with Dunnett’s test for multi-comparison correction.

Although we observed a negative correlation between NR2F6 expression and IFNγ signature genes in TCGA datasets relevant to breast, lung, and pancreatic cancers, inhibition of NR2F6 expression in cell lines obtained from these tumors did not increase their responsiveness to ICT, as seen in melanoma (Figures S3A-D), suggesting that factors present in melanoma but not in other tumors may be required for NR2F6 effects on the immune system.

### Tumor cell expression of NR2F6 controls T cell infiltration

To further define mechanisms underlying effects of tumor-intrinsic NR2F6 expression on anti-tumor immunity, we monitored the infiltration capacity of tumor infiltrating lymphocytes (TILs) into melanoma tumors engineered to lack NR2F6 expression. To this end, B16F10 tumor cells engineered to be NR2F6Nr2f6-deficient were injected into immune-competent mice, and tumors were collected 12 days later. Consistent with earlier observations, NR2F6 loss reduced tumor volume and weight compared with unmodified controls (Figure 3A). Notably, NR2F6-ablated tumors exhibited a 1.5-2 folds increase in infiltration of CD45^+^ immune cells, an increase more pronounced in the population of effector memory CD8^+^ T cells (2-2.5-fold increase of CD8^+^CD44^+^ cells; Figures 3B-D, Figure S4A). Increased CD8^+^ cell infiltration was accompanied by decreased B220^+^ B cell infiltration, but other immune cell types remained unchanged (Figure S4B). Consistent with increased CD8^+^ effector T cell infiltration, we observed a trend for increased IFNγ production and PD1 expression (Figure S4C-D), which coincided with increased activity of CD8^+^ T cells in NR2F6-deficient tumors. To confirm that CD8^+^ T cell infiltration mediates tumor regression seen after NR2F6 loss, we injected neutralizing antibodies for CD8^+^ T cells into mice inoculated with NR2F6-deficient B16F10 cells (Figure 3D). In contrast to mice treated with the IgG control antibodies, we observed no attenuation of melanoma growth or increased survival in mice harboring NR2F6-deficient B16F10 tumors after CD8^+^ T cell depletion (Figure 3E-F), suggesting that increased anti-tumor immunity seen in melanoma lacking NR2F6 expression is predominantly mediated by activity of tumor-infiltrating CD8^+^ T cells.

**Figure 3.**
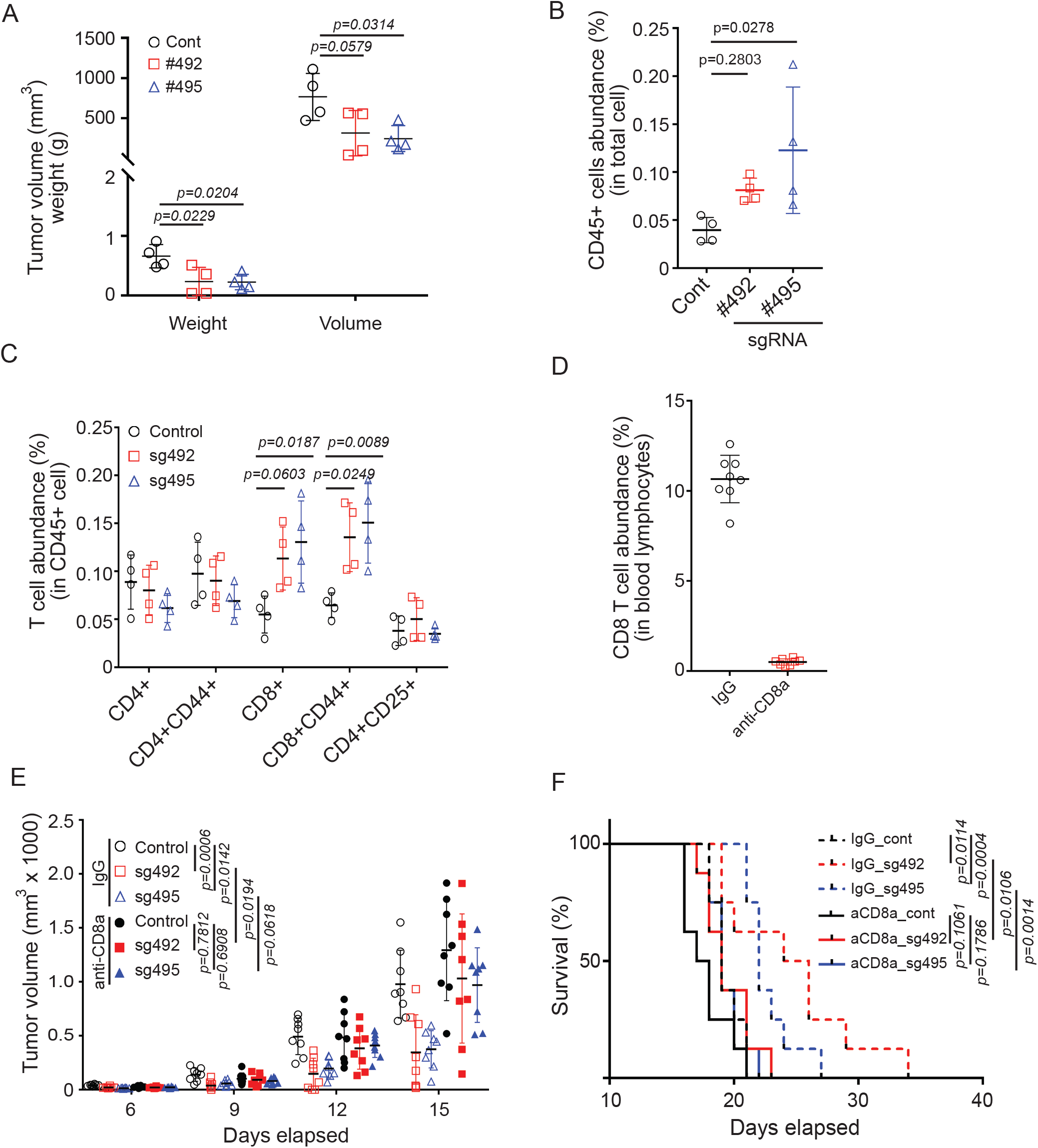
NR2F6 limits murine melanoma growth by controlling CD8 T cell infiltration. (A) Control and CRISPR-KO B16F10 cells were engrafted into C57BL/6 mice. Tumors were collected 12 days following their inoculation and assessed for weight and volume. (B) CD45^+^ cell abundance (expressed as a percentage) in cells within the singlet gate was assessed by FACS. (C) Abundance of T cell subtypes within all CD45^+^ cells were assessed by FACS. (D-F) Groups of mice were injected with control Ig or anti-CD8A antibodies, the latter to deplete CD8^+^ T cells. Then, CD8^+^ T cell abundance was assessed in blood samples collected 8 days later (D). Tumor growth (E) and overall animal survival (F) were monitored at indicated time points. Data are presented as means ± SD. Statistical significance was assessed by one-way ANOVA with Dunnett’s test (A, B, C), Student’s t-test (D), two-way ANOVA with Sidak’s test (E), or by long-rank test (F).

### RNAseq identifies NACC1 and FKBP10 as NR2F6 effectors

Given that tumor cell-intrinsic tumor effects of NR2F6 on anti-tumor immunity likely require NR2F6 transcriptional activity, we analyzed transcriptional changes seen following NR2F6 loss using RNAseq. To distinguish between changes in tumors vs. changes in infiltrating TME components, we performed RNAseq on (i) bulk tumor samples consisting of tumor and stromal components, (ii) MACS-sorted tumor cells (which lack stromal components), and (iii) cultured melanoma cells. Principal component analysis of RNAseq data indicated distinct gene expression patterns in each sample (Figure S5A-B). Of note, one of the two sgRNAs used for NR2F6 knockout (KO), sg495, resulted in a better separation of tumor from control samples (Figure S5B), and thus we focused on this sample set. In bulk tumors, sorted tumor cells, and cultured cells, NR2F6 KO resulted in differential upregulation of 361, 245, and 106 genes and downregulation of 57, 102, and 310 genes, respectively (Figures 4A, S5C). Consistent with tumor growth inhibition seen upon NR2F6 loss, both pathway and upstream regulator analyses of differentially expressed genes (DEGs) identified activation of anti-tumor immune signaling only in in vivo (bulk tumor and sorted tumor cells) samples. Among pathways upregulated by NR2F6 KO were inflammation, T cell receptor, and Th1 and IFN signaling, with common upstream pathways responsive to LPS and IFNγ stimuli as to STAT1 signaling (Figures 4B-C). By contrast, we observed no activation of immune-related pathways in cultured melanoma cell samples (Figures 4B-C), suggesting the importance of an in vivo setting in controlling immune cell components.

**Figure 4.**
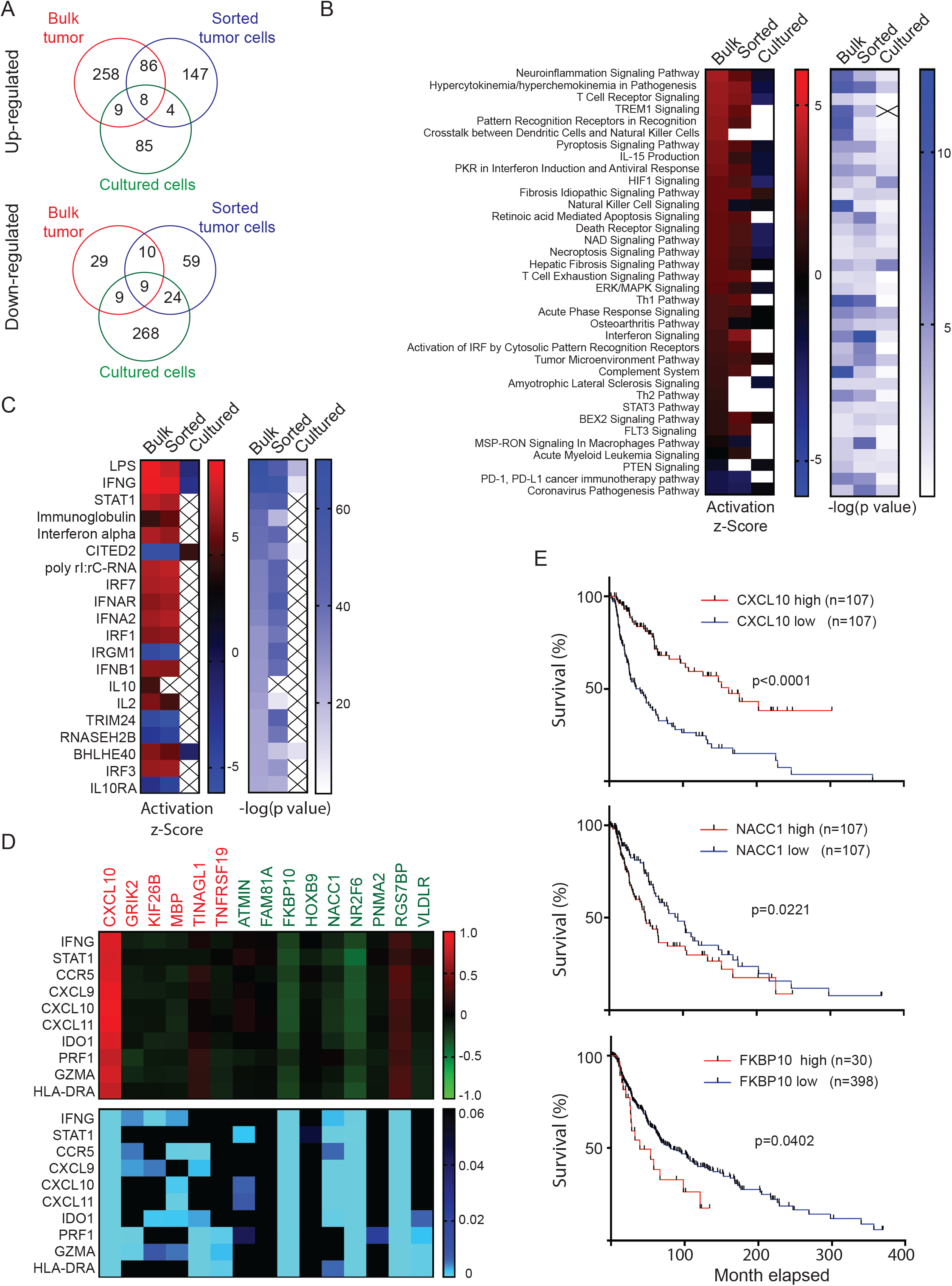
RNAseq identifies NACC1 and FKBP10 as NR2F6 effectors. (A) Differentially expressed genes (DEGs) between control and CRISPR-KO B16F10 were identified based on RNAseq analysis of bulk tumors, MACS-sorted tumor cells or cultured cells. Unique or common genes upregulated (upper) or downregulated (lower) were plotted in Venn diagrams. (B-C) Activated or repressed pathways (B) and upstream regulators (C) of identified DEGs were analyzed using Ingenuity Pathway Analysis (IPA). (D) Heatmaps show the Spearman correlation coefficient (upper) and corresponding p values (lower) in the comparison of expression of 15 genes commonly up- or down-regulated in RNAseq data from the three samples described in (A). X- and Y-axes indicate up/down-regulated genes and 10 IFNγ signature genes, respectively. CXCL10, FKBP10 and NACC1 were significantly correlated with NR2F6 and IFNγ signature. (E) Overall survival of melanoma patients with low or high expression of CXCL10, FKBP10 or NACC1 in melanoma specimens, as assessed by Kaplan-Meier analysis. Statistical significance was assessed by long-rank test.

Next, to identify downstream effectors for NR2F6, we assessed an additional 17 genes up- or down-regulated in NR2F6 KO samples, including bulk tumors, sorted, and cultured cells, for (i) possible association with IFNγ signature genes and (ii) correlation with patient survival (Figure 4D-E). Of 9 genes downregulated by NR2F6 KO, NACC1 and FKBP10 were negatively correlated with the IFNγ-signature, and high NACC1 and FKBP10 expression was associated with poor overall survival, patterns resembling changes seen after tumor-intrinsic NR2F6 loss. Notably, NACC1 and NR2F6 expression was high in all human melanoma lines and in patient tissue samples analyzed (Figure S5D).

### NACC1 and FKBP10 loss phenocopies enhanced anti-tumor immunity seen upon NR2F6 loss

To further analyze activity of NACC1 and FKBP10 as NR2F6 effectors, we monitored NACC1 and FKBP10 expression levels in NR2F6 KD or KO mouse and human melanoma lines. Both NACC1 and FKBP10 expression was downregulated in mouse melanoma cells subjected to NR2F6 KD or KO, consistent with our RNAseq analysis (Figure 5A-B, S6A). To confirm these observations, we asked whether loss of NACC1 and/or FKBP10 would phenocopy effects of NR2F6 loss, attenuating tumor growth. To do so, we established B16F10 melanoma cells deficient in either NACC1 or FKBP10 that were then injected into immune-competent and -compromised mice and monitored for tumor size. NACC1 or FKBP10 depletion attenuated tumor growth (33.2 ∼ 44.7% reduction) in immune-competent but not immune-compromised mice (Figure 5C-H), resembling effects seen in NR2F6-depleted melanoma, while we did not observe overt changes in cell growth in cultured cells (Figure S6C-D). These observations suggest that NACC1 or FKBP10 loss suppresses tumor growth possibly by augmenting anti-tumor immunity. Notably, combined loss of both NACC1 and FKBP10 further limited tumor growth (52.83% reduction) in immune-competent but not - compromised mice, again without altering cell growth *in vitro* (Figure 6A-D).

**Figure 5.**
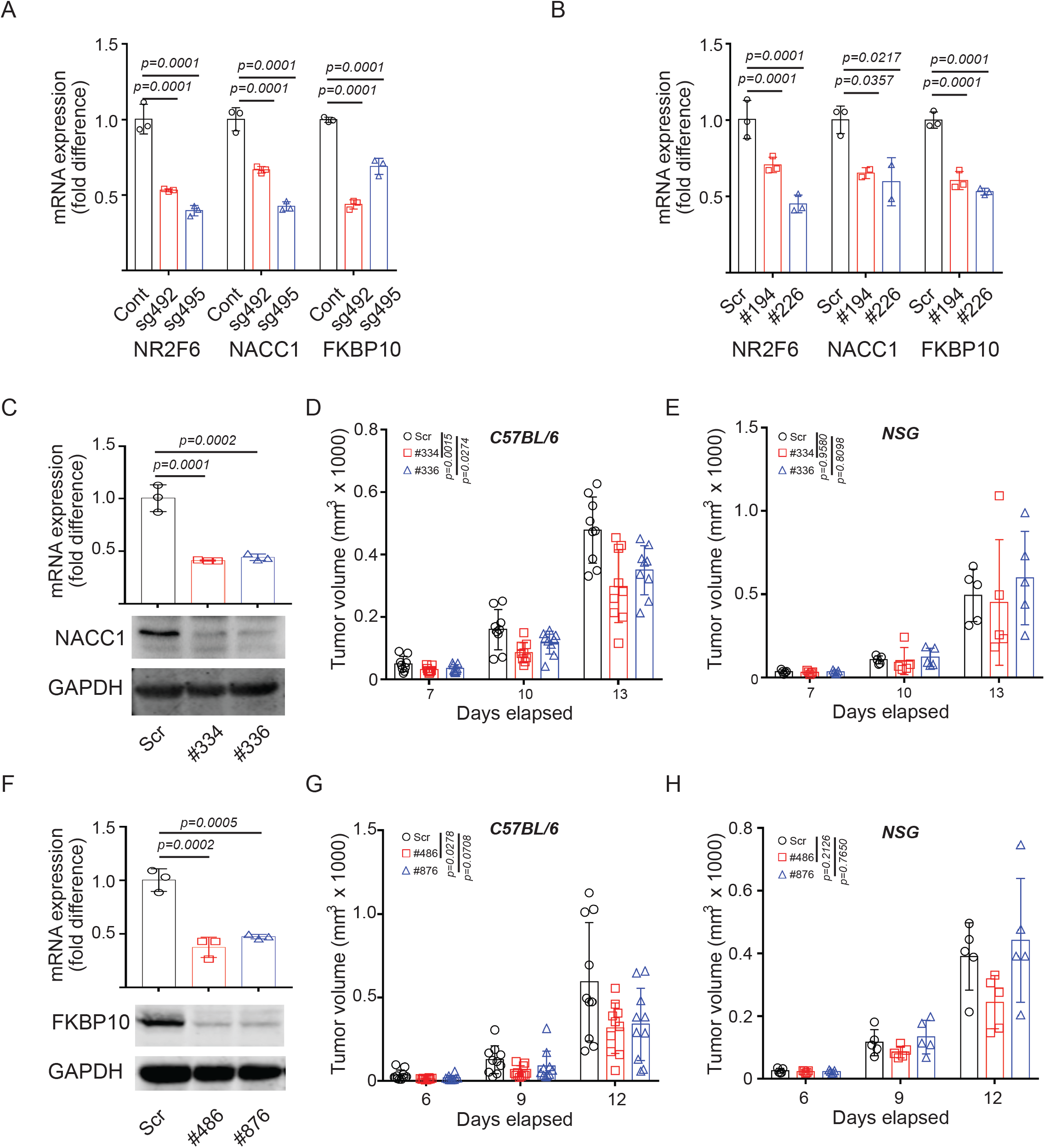
Loss of NACC1 or FKBP10 attenuated tumor growth in mice with intact immune system. (A-B) The expressions of indicated genes were assessed in B16F10 KO (CRISPR-based) (A) or KD (shRNA based) (B) cells by qPCR. (C) B16F10 cells were transduced with scrambled (Scr) and two shRNAs (#334 and #336) against NACC1. mRNA and protein expression were assessed by qPCR and immunoblotting. (D-E) Then, cells were inoculated to C57BL/6 (D) or NSG (E). Tumor volumes were monitored at indicated time points. (F-H) As in (C-E), B16F10 cells were transduced with scrambled and two (#486, #876) shRNAs against FKBP10. FKBP10 expressions were assessed (F). Tumor growth of transduced cells were monitored in C57BL/6 (G) or NSG (H) mice. Data are presented as means ± SD. Statistical significance was assessed by one-way ANOVA with Dunnett’s test (A, B, C, F), or two-way ANOVA with Dunnett’s test (D, E, G, H).

**Figure 6.**
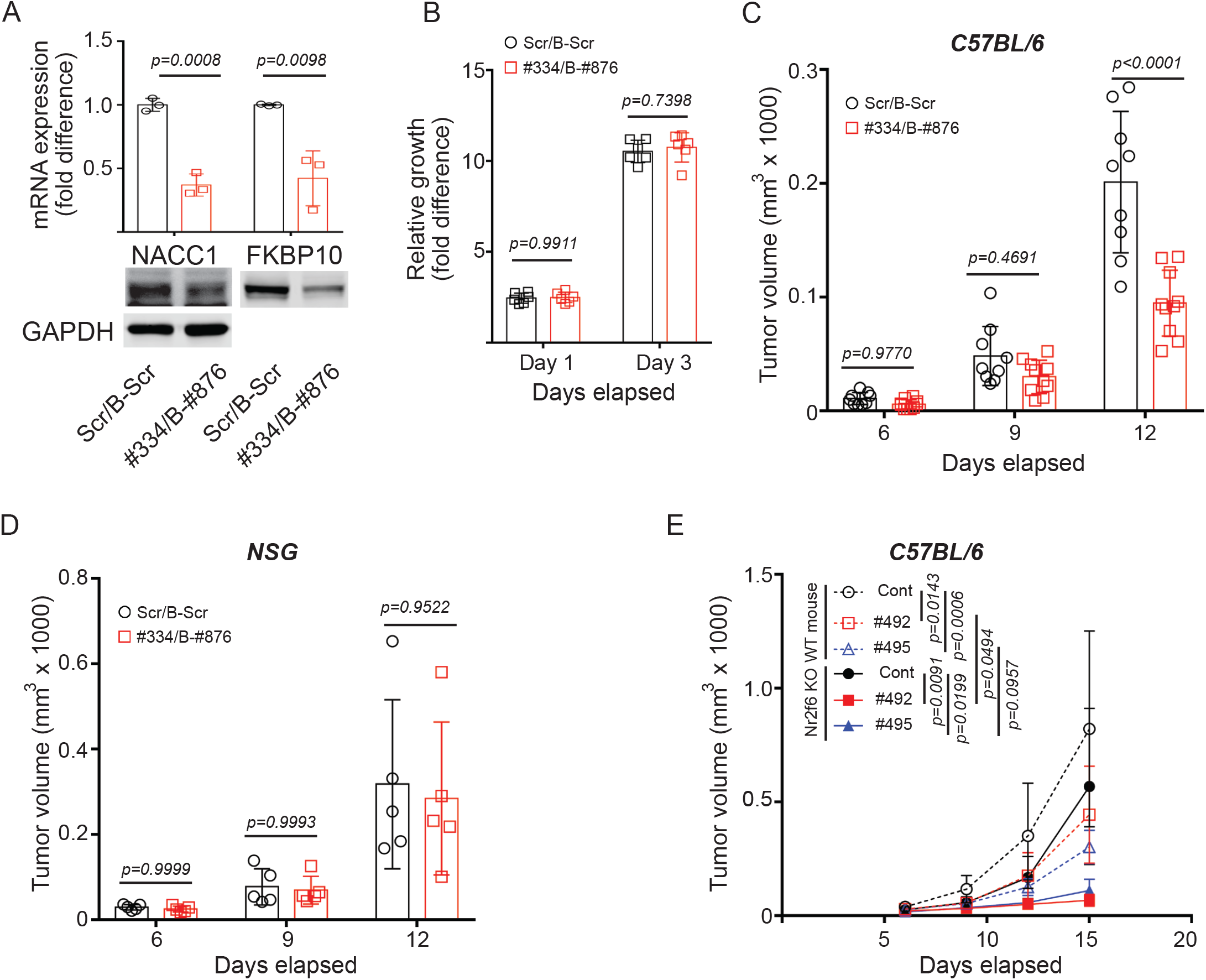
Loss of NACC1 and FKBP10 phenocopies NR2F6 loss; combination of intrinsic and extrinsic NR2F6 loss augments degree of melanoma inhibition. (A) B16F10 cells were transduced with scrambled [Scr (puromycin resistant), B-Scr (blasticidin resistant)] and two shRNAs (#334 for NACC1 KD and #876 for FKBP10 KD). mRNA and protein expression were assessed by qPCR and immunoblotting. (B) Growth of transduced cells *in vitro* was assessed using CellTiter-Glo. (C-D) The transduced cells were inoculated to C57BL/6 (C) or NSG (D) mice. Tumor growth was monitored at indicated time points. (E) B16F10 control cells (NR2F6 WT) or NR2F6 KO cells (#492 and #495) were inoculated to C57BL/6 control or NR2F6 KO mice. Tumor volumes were monitored at indicated time points. Data are presented as means ± SD. Statistical significance was assessed by student’s t-test (A, B, C, D), or two-way ANOVA with Tukey’s test (E).

### Anti-tumor immunity seen following loss of tumor intrinsic NR2F6 is augmented by systemic NR2F6 loss

The ability of tumor extrinsic NR2F6 expression to block anti-tumor immunity was previously demonstrated in genetically ablated NR2F6 KO mice, which exhibit suppressed anti-tumor immunity (Hermann-Kleiter *et al*., 2015; Klepsch *et al*., 2018). Thus, we asked whether the tumor-intrinsic function of NR2F6 loss in promoting anti-tumor immunity demonstrated here would be augmented in mice globally deficient in NR2F6. To this end, we inoculated both WT and NR2F6 KO mice with B16F10 cells that either harbored or lacked NR2F6. Compared to either NR2F6 depletion in tumor cells only (45.85 ∼ 62.95% reduction) or systemic NR2F6 loss only (30.77% reduction), the combined depletion of NR2F6 in both tumors and systemically, that is, in all TME cells, further decreased melanoma growth (86.37 ∼ 91.69% reduction) (Figure 6E). These findings suggest that both tumor-intrinsic and -extrinsic NR2F6 expression contributes to anti-tumor immunity and to the degree of tumor growth inhibition.

## Discussion

Deregulated NR expression has been documented in several tumor types, as in number of immune cell types. Here, we identify a previously undisclosed role for the orphan nuclear receptor NR2F6 in tumor-intrinsic control of immune evasion by melanoma cells, a function highly relevant to ICT effectiveness. NR2F6 expression in tumor cells coincided with that of IFNγ signature genes and with overall melanoma patient survival, which formed the basis for our focus on NR2F6 among a group of >40 candidates. Accordingly, others previously identified that NR2F6 was often deregulated and associated with therapy resistance and disease recurrence in other cancer types, including head and neck squamous cell carcinoma and ovarian cancers (Klapper et al., 2020; Li *et al*., 2019). Our findings establish NR2F6’s tumor-intrinsic function and identify NACC1 and FKBP10 as its downstream transcriptional targets, which mediate control of anti-tumor immunity. The KD of either NR2F6, NACC1, or FKBP10 in melanoma cells led to tumor growth inhibition in immune-competent but not -incompetent mice. Albeit the underlying mechanism remain elusive, this data supports the notion that NACC1 and FKBP10 are NR2F6 effectors which regulate tumor evasion by the immune system. Notably, while NR2F6 KD effectively induced immune cell infiltration, primarily by CD8^+^ T cells, which limited tumor growth, NR2F6 overexpression did not enhance tumor growth, suggesting either that co-factors (which were not overexpressed) are needed for NR2F6 function or that tumor intrinsic roles of NR2F6 are dependent on the availability of tumor extrinsic factors. The latter is plausible, given that a tumor-extrinsic role for NR2F6 in controlling anti-tumor immunity in two distinct autochthonous tumor models shown in mice genetically (globally) deficient in NR2F6 (Hermann-Kleiter *et al*., 2015; Klepsch *et al*., 2018).

Importantly, previous studies employing subcutaneous NR2F6 expressing wild-type tumor cell graft models suggest that NR2F6 can regulate immune system activities independently of its tumor expression, suggesting a NR2F6 function in the TME, and more specifically when employing acute NR2F6 gene silencing prior adoptive transfer in CD8+ T cells (Klepsch *et al*., 2020). Along these lines, as reported previously, genetic ablation of NR2F6 in mice potentiated anti-tumor PDL1/PD1 ICT (Klepsch *et al*., 2018).

Yet, here, combined ablation of NR2F6 in both melanoma cells and globally had further increased the degree of tumor inhibition, suggesting that distinct but complementing factors contribute to enhancement of anti-tumor immunity. Future studies should address potential tumor-extrinsic factors that regulate the degree of anti-tumor immunity. Among candidates are non-immune stromal components, which express higher levels of NR2F6 than do immune cell subtypes (supplementary figure S1B-C). A role for stroma in control of immune cell infiltration and activation has been established (Barrett and Pure, 2020; Nagl et al., 2020); thus NR2F6 controls stromal cell activities in a way that recruits or activates CD8^+^ T cells and promotes their infiltration of tumors. Accordingly, IFNγ reportedly plays an important role in NR2F6-dependent enhancement of CD8^+^ T cell memory during early immune response to bacterial infection (Jakic *et al*., 2021).

How does KD of NR2F6 or its effectors NACC1 and FKBP10 enhance recruitment of CD8^+^ T cells to limit tumor growth? NACC1 (aka NAC1) protein reportedly serves as a bridge between MAVS and TBK1 proteins to induce anti-viral signaling, which could activate innate immunity (Xia et al., 2019), although NACC1 has not been previously linked to immune evasion. Likewise, FKBP10 activation is proposed to attenuate anti-tumor immunity in a colorectal cancer model (Chen et al., 2022), a finding consistent with activation of anti-tumor immunity reported here after FBBP10 KD.

Could NR2F6 serve as a target for therapy? This strategy has been suggested by the Baier group, who demonstrated the importance of NR2F6 as an intracellular immune checkpoint in effector T cells (Klepsch et al., 2016) and later showed that mice genetically ablated of NR2F6 and inoculated with tumor cells exhibited decreased tumor growth (Hermann-Kleiter *et al*., 2015; Klepsch *et al*., 2018). Here we add an important component to this equation by demonstrating that tumor-intrinsic NR2F6 expression is equally relevant, and that combining tumor-intrinsic and -extrinsic ablation of NR2F6 synergizes to block tumor growth. This finding justifies development of NR2F6 inhibitors that could be administered systemically to target both tumors and the TME. Initial efforts to identify such compounds have been reported (Smith et al., 2022) and should gain more traction given findings reported here.

## Materials and Methods

### Experimental models

All studies conducted in mice were approved by the Institutional Animal Care and Use Committee of Sanford Burnham Prebys Medical Discovery Institute (AUF#21-032). Murine melanoma (B16F10, YUMM1.7, and YUMMER1.7), and breast (4T-1), lung (LLC), and pancreatic ductal adenocarcinoma (KPC) cancer lines were injected subcutaneously (2.0 × 10^5^ cells of B16F10 or YUMM1.7; 4.0 × 10^5^ cells of YUMMER1.7; 1.0 × 10^6^ cells of 4T-1; 1.5 × 10^6^ cells of LLC) into the lower right flank of 6-8-week-old male C57BL/6 (B16F10, YUMM1.7, YUMMER1.7, LLC), female Balb/C (4T-1), Nod-Scid-Gamma (NSG) (B16F10, YUMM1.7) or NR2F6 knockout (Klepsch *et al*., 2018) mice. Tumor growth was monitored weekly using calipers. At indicated time points, tumors were collected, weighed, and assessed for immune phenotypes using flow cytometry. To assess tumor response to immune checkpoint antibodies, mice were grafted with B16F10 (2.0 × 10^5^ cells, s.c.) cells and treated with 200 µg/mouse anti-CD279 (PD-1) [RMP1-14 (BE0146, BioXcell)]. Antibodies were injected (i.p.) 3 - 5 times (every 3 days starting at the indicated date). To assess percent survival of animals, mice bearing tumors exceeding 2,000 mm^3^ were defined as “dead”.

### Analysis of cancer gene expression and patients’ clinical outcomes

To correlate expression of 48 NRs and IFNγ-signature genes (Ayers *et al*., 2017), mRNA expression data (Z-scores relative to diploid samples) of those genes in TCGA datasets (bulk RNAseq) were downloaded from cBioPortal (Cerami et al., 2012; Gao et al., 2013). Spearman correlation coefficients between expression of NRs and IFNγ-signature genes were analyzed. A corresponding heatmap based on correlation coefficient and p value data was plotted using Prism7.0 software (GraphPad).

The IFNγ-index for each patient sample was calculated as an average rank percentile of 10 IFNγ signature genes in a total of 443 samples. Based on that value, groups of patients with an index >75% or <25% were selected for further assessment of correlation with overall survival which was obtained from cBioPortal. The correlation of IFNγ-index with patient overall survival was assessed by Kaplan-Meier survival analysis using Prism7.0 (GraphPad). Likewise, the correlation of patient overall survival with NR expression was assessed. Briefly, 2 groups of patients, 25^th^ and 75^th^ quartiles in NR expression, were compared by Kaplan-Meier survival analysis. NRs with a significant p-value in the log-rank (Mantel-Cox) test were plotted in the heatmap as exhibiting either favorable or unfavorable overall survival.

Patient response to ICT and corresponding NR expression in malignant cells was plotted using published data and analysis (Single Cell portal, (Jerby-Arnon *et al*., 2018). To assess NR expression in each cell type found in melanoma tumors, we downloaded single cell RNAseq data (GSE72056, (Tirosh et al., 2016). The percent expression of each NR different cell types was calculated by dividing the number of cells expressing NR by the total cell count.

### RNAseq analysis

For gene expression analysis, RNA samples were prepared from bulk tumors, MACS-sorted tumor cells and cultured cells. To purify tumor cells from bulk tumor tissue, collected B16F10 tumors were minced, chopped and incubated in Collagenase D solution [0.1 % (w/v) Collagenase D, 0.5 % (w/v) BSA, 100 µg/ml DNase in PBS] for 1 h at 30 **°**C. A single cell suspension was obtained by using a cell strainer (70µm, Falcon). Tumor cells were purified by depletion of stromal cells using a Tumor Cell Isolation kit (Miltenyi Biotech). Purity of isolated tumor cells was validated by FACS using CD45 antibody. RNA from bulk tumors, MACS-sorted tumor cells, and cultured cells was purified using a GenElute total RNA purification kit (Sigma-Aldrich).

For library construction, PolyA RNA was isolated using NEBNext® Poly(A) mRNA Magnetic Isolation Module, and bar-coded libraries were constructed using the NEBNext® Ultra™ Directional RNA Library Prep Kit for Illumina® (NEB, Ipswich MA). Libraries were pooled and sequenced single-end (1×75) on an Illumina NextSeq 500 using the High output V2 kit (Illumina Inc., San Diego CA) at a sequencing depth of 23-31 million reads. Raw reads were trimmed to remove llumina Truseq adapters and polyA/polyT sequences using Cutadapt version 2 (Martin, M, 2011). Reads were then aligned to mouse genome version mm10 and Ensembl gene annotations version 84 using STAR version 2.7.0d_0221 (Dobin et al., 2013) and alignment parameters from ENCODE long RNA-seq pipeline (https://github.com/ENCODE-DCC/long-rna-seq-pipeline). We obtained gene level estimated counts and transcripts per million (TPM) using RSEM version 1.3.1 (Li and Dewey, 2011). FastQC version 0.11.5 (https://www.bioinformatics.babraham.ac.uk/projects/fastqc/) and MultiQC version 1.8 (Ewels et al., 2016) were used to assess the quality of trimmed reads and alignment to genome/transcriptome. We removed low expressed genes from downstream analysis by selecting those with RSEM estimated counts that are equal or greater than 5 times the total number of samples. Differential expression comparisons were performed using Wald test implemented in DESeq2 version 1.22.2 (Love et al., 2014). Genes with Benjamini-Hochberg corrected p-value < 0.05 and fold change ≥ 1.5, or ≤ -1.5, were identified as differentially expressed. Pathway analysis was performed using Ingenuity Pathway Analysis (Qiagen, Redwood City, USA)

### Cloning DNA constructs and mutagenesis

DNA plasmids were constructed using the pLX302 and pLX304 Gateway system (Addgene, #25890). Briefly, PCR-amplified mouse mouse *Nr2f6, Nr1h3*, and *Nr3c1* cDNAs were cloned into the lentiviral Gateway Vector using LR clonase II and a pENTR-D-TOPO cloning kit (Thermo Fisher Scientific). A mutant form of Nr2f6 (C112S) incapable of binding DNA was generated by introducing point mutations into the pENTR-Nr2f6 construct using a QuikChange II XL Site Directed Mutagenesis Kit (Agilent), and the inserts were subsequently cloned into pLX304 (lentiviral) expression plasmids. shRNA clones harboring a blasticidin-resistance gene were generated by cloning validated oligonucleotides into the EcoRI/AgeI sites of pLKO.1-Blast (Addgene, #26655). Gene-specific shRNA lentiviral vectors with a pLKO.1 backbone were purchased from Sigma-Aldrich.

### Production and infection of viral particles

Lentiviral particles were prepared using standard protocols. Briefly, HEK293T cells were transfected with lentiviral plasmid and the second-generation packaging plasmids psPAX2 (Addgene, #12260) and pMD2.G (Addgene, #12259) using Calfectin (SignaGen) or JetPrime (Polyplus). Viral supernatants were collected 48 hr later, filtered using a syringe filter (0.45 µm pore size) and concentrated by centrifugation (13, 000 x g for 2hr). Titrated viral particles and polybrene (8 µg/ml, Sigma) were applied to the culture (2 × 10^5^ cells/well in 6-well plates) and subjected to spinoculation (1,500 x g for 30 min) to infect melanoma cells. To establish stably-transduced cells, efficiently-infected cells were selected in cultures containing either puromycin (InvivoGen, 1 ∼ 1.5 µg/ml) or blasticidin (InvivoGen, 5 ∼ 10 µg/ml), as appropriate.

### Gene Silencing

To knock down genes, pLKO.1clones and esiRNA for respective genes were purchased (Sigma-Aldrich). Specific pLKO.1 clones for each gene were as follows: Nr2f6 (TRCN0000026194, TRCN0000026226, TRCN0000033661, TRCN0000033662, TRCN0000033663), Nacc1 (TRCN0000071334, TRCN0000071336) and Fkbp10 (TRCN0000339486, TRCN0000111876). For double knockdowns, oligonucleotides of the same sequence (TRCN0000111876) were cloned into pLKO.1-blast (Addgene #26655). Cells transduced were selected in culture containing puromycin (InvivoGen) or blasticidin (Gibco).

To knock out Nr2f6 using CRISPR, Nr2f6-specific sgRNAs (Thermo Fisher Scientific, CRISPR553495_SGM, CRISPR553492_SGM) were labeled with Cy3 using a Label IT kit (Mirus). B16F10 cells were transfected with labeled sgRNAs and Cas9 protein (Thermo Fisher Scientific) using CRISPRMax Cas9 transfection reagent (Thermo Fisher Scientific). Cells were collected after 24 h of culture and subjected to FACS-sorting to isolate Cy3^+^ cells. Nr2f6 KO was validated by immunoblotting and subsequent sequencing of regions targeted by sgRNAs.

### Assessment of cell growth in culture

Cell growth in culture was measured by assessing ATP content of viable cells using CellTiter-Glo (Promega). Briefly, 2,000-2,500 cells were placed into 96-well plates with clear bottoms (Nunc), CellTiter working solution was added to wells, and luminescence was measured with a Clariostar microplate reader. Luminescence at days 1 and 3 was calculated as a fold-difference relative to luminescence at day 0.

### Immunoblotting

Immunoblotting samples were processed using standard protocols with slight modifications. Briefly, cells were lysed by incubation in RIPA buffer with 0.1% SDS (sodium dodecylsulfate) [50mM Tris-HCl, pH7.4, 1% (v/v) NP40, 0.1% (w/v) sodium deoxycholate, 0.1% (w/v) SDS, 150mM NaCl, 1mM EDTA, and a protease/phosphatase inhibitor cocktail (Thermo Fisher Scientific)] and 3 freeze-thaw cycles. Tumor tissues were lysed by incubation in RIPA buffer with 0.5% SDS [50mM Tris-HCl, pH7.4, 1% (v/v) NP40, 0.1% (w/v) sodium deoxycholate, 0.5% (w/v) SDS, 150mM NaCl, 1mM EDTA, and a protease/phosphatase inhibitor cocktail (Thermo Fisher Scientific)] followed by homogenization using a Tissuemiser (Fisher Scientific). Lysates were boiled in Laemmli buffer before separation on SDS-PAGE and then transferred to a PVDF membrane. Membranes were incubated with blocking solution [TBS (Tris-buffered Saline); 10 mM Tris-HCl, pH 8.0, 150 mM NaCl)] containing 0.1% Tween 20 and 5% nonfat milk followed by incubation with appropriate primary antibodies overnight at 4 °C. Membranes were washed with TBS and incubated 1 hr at room temperature with secondary antibody [Alexa 680-conjugated goat anti-rabbit, goat anti-mouse, donkey anti-goat (Life Technologies) or IRDye 800-conjugated goat anti-mouse (Rockland Immunochemicals)], or HRP-conjugated anti-mouse or anti-rabbit IgG antibodies (Cell Signaling). Bands on blots incubated with fluorescent antibodies were visualized and quantified using an Odyssey Infrared Imaging System (LiCoR Biosciences). Bands with HRP activity were visualized using a ChemiDoc Imaging system (Bio-Rad) after incubating blots with West Pico plus Chemiluminescent substrate (Thermo scientific) or Immobilon Forte Western HRP substrate (Millipore). The following antibodies were used to detect respective proteins: NR2F6 (60117-1-Ig, 60117-2-Ig, Proteintech), NAC1 (#4183, #4420, Cell Signaling), FKBP10 (12172-AP, Proteintech), and V5-tag (7/4, Biolegend).

### RNA extraction and quantitative PCR (QPCR)

Total RNA was purified from cells with GenElute (Sigma-Aldrich), a PureLink RNA kit (Invitrogen) or a Quick-RNA kit (Zymo Research). Purified RNA was reverse-transcribed using a high-capacity cDNA synthesis kit (Applied Biosystems). qPCR was carried out with a CFX Connect Real-Time PCR Detection System (Bio-Rad) using SsoAdvanced Universal SYBR Green Supermix (Bio-Rad). Sequences of primers used were as follows:

18S rRNA (internal control for normalization):

5’-GTAACCCGTTGAACCCCATT-3’, 5’-CCATCCAATCGGTAGTAGCG-3’

mouse Nr2f6:

5’-GAGGACGATTCGGCGTCAC-3’, 5’-GTAATGCTTTCCACTGGACTTGT-3’

human NR2F6

5’-GAGCGGCAAGCATTACGGT-3’, 5’-GGCAGGTGTAGCTGAGGTT-3’

mouse Nacc1:

5’-GCGGCTACAGGGACTATACTG-3’, 5’-CCGGAAGTAAGAGCTACTAGCG-3’

Human NACC1:

5’-CTGGCTCCTACCACAATGAGG-3’, 5’-TGGCCGACGTTCATCATGC-3’

mouse Fkbp10:

5’-TACTGCCGTTGCTGTTGCTT-3’, 5’-GGGATGTGGTATCTCTCGATGAC-3’

Human FKBP10:

5’-TACAGTAAGGGCGGCACTTAT-3’, 5’-GAGGACGTGAAAGACCAGCG-3’

mouse Cxcl10:

5’-CCAAGTGCTGCCGTCATTTTC-3’, 5’-GGCTCGCAGGGATGATTTCAA-3

### Immune phenotyping of tumors using flow cytometry analysis

To assess the immune phenotypes of tumor-infiltrating lymphocytes (TILs), B16F10 tumors were collected at indicated times and then chopped and incubated in Collagenase D solution [0.1 % (w/v) Collagenase D, 0.5 % (w/v) BSA, 100 µg/ml DNase in PBS] for 1 h at 30 **°**C. A single cell suspension was obtained using a cell strainer (70µm, Falcon). Total cells were counted and a fraction (2 × 10^6^) of cells in FACS staining buffer (phosphate-buffered saline, pH 7.4, containing 1 % FBS) was treated with the following sets of antibodies (1:200 dilution): cocktail 1 [CD45.2 (AF700), CD8 (APC), CD4 (BV605), CD44 (APC/Cy7), CD25 (FITC) and purified CD16/32], cocktail 2 [CD45.2 (AF700), MHCII (PB), CD11C (APC), CD11b (APC/Cy7), GR1 (PE), F4/80 (FITC), NK1.1

(BV605), B220 (PE/Cy7) and purified CD16/32], and cocktail 3 [CD45.2 (AF700), CD8 (FITC), CD4 (BV605), CTLA4 (PERCP/Cy5.5), LAG3 (PE), PD-1 (APC), TIM-3 (PE/Cy7)] for 20 min at 4 **°**C. All antibodies were from Biolegend. Stained cells were fixed in 1% formaldehyde (Sigma) in PBS (pH 7.4) for 15 min at 4 **°**C and analyzed with BD LSR Fortessa (BD Biosciences) flow cytometry. To assess intracellular cytokines in infiltrated CD4^+^ and CD8^+^ cells, a fraction (2 × 10^6^) of cells prepared from tumors was stimulated with PMA (10 ng/ml)/ionomycin (0.5 µg/ml)/BFA (1 µg/ml) for 16 hr. Cells were stained with a cocktail of antibodies for surface markers [CD45.2 (AF700), CD4 (BV605), and CD8 (APC)] followed by staining with intracellular cytokine antibodies [IFNγ (APC), TNFα (FITC) and IL-2(PE)]. The abundance of each cell type was calculated as a percentage of CD45^+^ cells.

### CD8 T cell depletion

To deplete CD8^+^ T cells, mice were treated with anti-CD8^+^ antibody [2.43 (BE0061, BioXcell), while controls were treated with 200 µg/mouse IgG [rat IgG2b (BE0090, BioXcell)]. Antibodies were injected (i.p.) every 3 days starting one day prior to tumor cell inoculation. The efficiency of depletion was assessed using flow cytometry of blood samples collected at day 8 after tumor inoculation.

### Statistical Analysis

Statistical analyses were performed using Prism software (version 7.00, GraphPad). For comparison of means of two groups with normal (or approximately normal) distributions, an unpaired t-test was applied. In multiple t-tests between two groups, adjusted p values were computed using the Holm-Sidak method. To compare means between >2 groups, we used one-way analysis of variance (ANOVA) with multiple comparison corrections (Dunnett’s test). For animal experiments, we used two-way ANOVA (time and treatment) with Dunnett’s, Tukey’s, or Sidak’s multiple comparison test. For Kaplan-Meier plots to compare overall survival, we used a log-rank (Mantel-Cox) test to determine the significance of differences between groups.

## Acknowledgments

We thank members of the Ronai lab for continued discussions. Support by grant R35CA197465 to ZAR is gratefully acknowledged. Shared resources that were part of this study (animal, cell sorting and bioinformatics) are supported by P30CA030199, Cancer Center Support Grant.

## Author contributions

HK and ZAR designed; HK performed the studies. RM performed bioinformatic analyses. GB and VK generated and provided the Nr2f6 KO mice. ZAR and HK wrote the manuscript.

## Supplemental Figures

**Supplementary figure S1.**
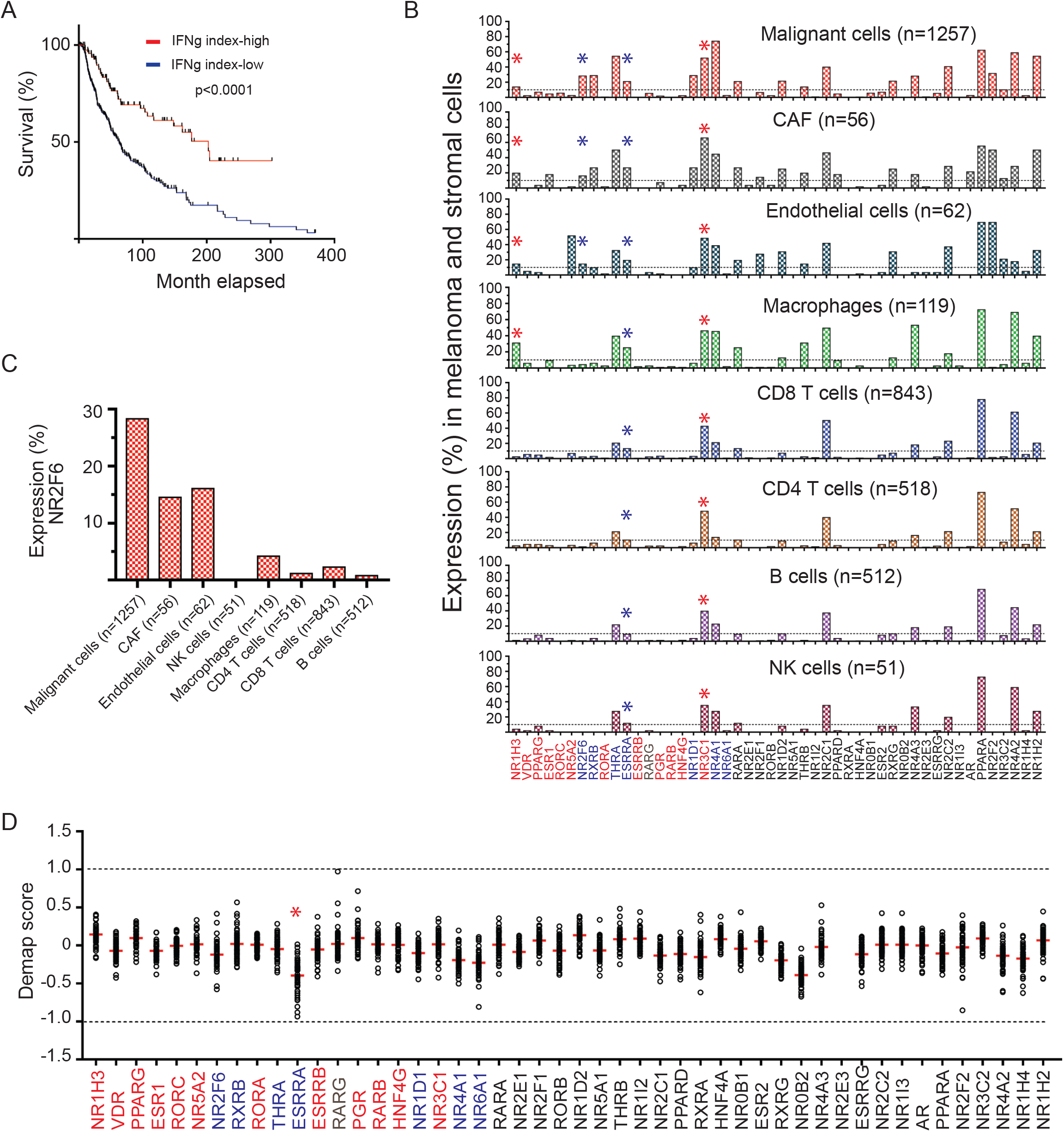
Identification of NRs that are associated with anti-tumor immunity. (A) Overall survival of melanoma patients with a high- or low-IFNγ signature (see Methods), was determined based on Kaplan-Meier analysis. (B) The percent expression of NRs in indicated cell types was analyzed by scRNAseq of human melanoma tissues (Tirosh *et al*., 2016). (C) The percent expression of NR2F6 in the indicated cell types was assessed using the same dataset. (D) DepMap (https://depmap.org/portal/) scores indicate effects on cell growth following KO of NRs in human melanoma lines. Each data point indicates a different line.

**Supplementary figure S2.**
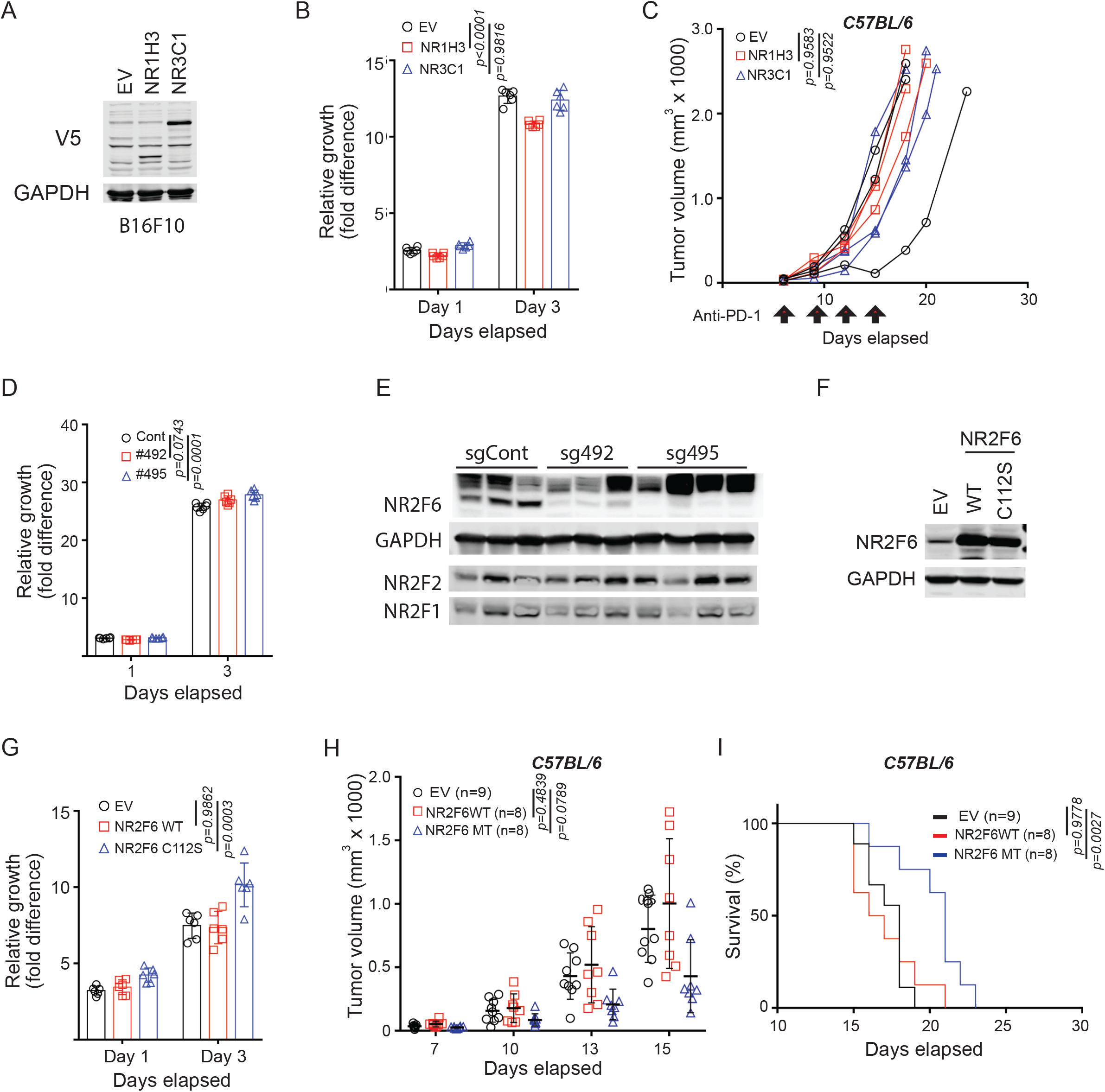
Validation of selected NRs in the control of anti-tumor immunity. (A) B16F10 cells that stably express V5-tagged NR1H3 or NR3C1 were established by transduction with the corresponding lentivirus constructs. Cell lysates were analyzed by immunoblotting with an antibody to V5-tag. (B) Growth of cultured cells established in (A) was assessed *in vitro* using CellTiter-Glo. Relative difference in luminescence values at days 1 and 3 were calculated based on level of luminescence at day 0, which was defined arbitrarily as 1. (C) Cells were inoculated in C57BL/6 mice, which were treated with anti-PD-1 antibody (RMP1-14) at days 6, 9, 12, and 15 (arrows). Tumor volumes were monitored at the indicated time points. (D) Growth of control and CRISPR-KO B16F10 cells (Figure 2F), as assessed using CellTiter-Glo. (E) Lysates established from mouse tumors (Figure 2G) were analyzed by immunoblotting of indicated proteins. (F) B16F10 cells stably overexpressing either WT or C112S (DNA-binding mutant) NR2F6 constructs were established by transduction of the corresponding lentiviruses, followed by immunoblotting of cell lysates for the indicated proteins. (G) The growth of the transduced cells *in vitro* was assessed using CellTiter-Glo, as outlined in (B). (H-J) The transduced cells were inoculated in C57BL/6 mice followed by monitoring tumor volumes and mouse survival at the indicated time points. Data are presented as means ± SD. Statistical significance was assessed by one-way ANOVA with Dunnett’s test (B, D, G), two-way ANOVA with Dunnett’s test (C, H), or by long-rank test (I).

**Supplementary figure S3.**
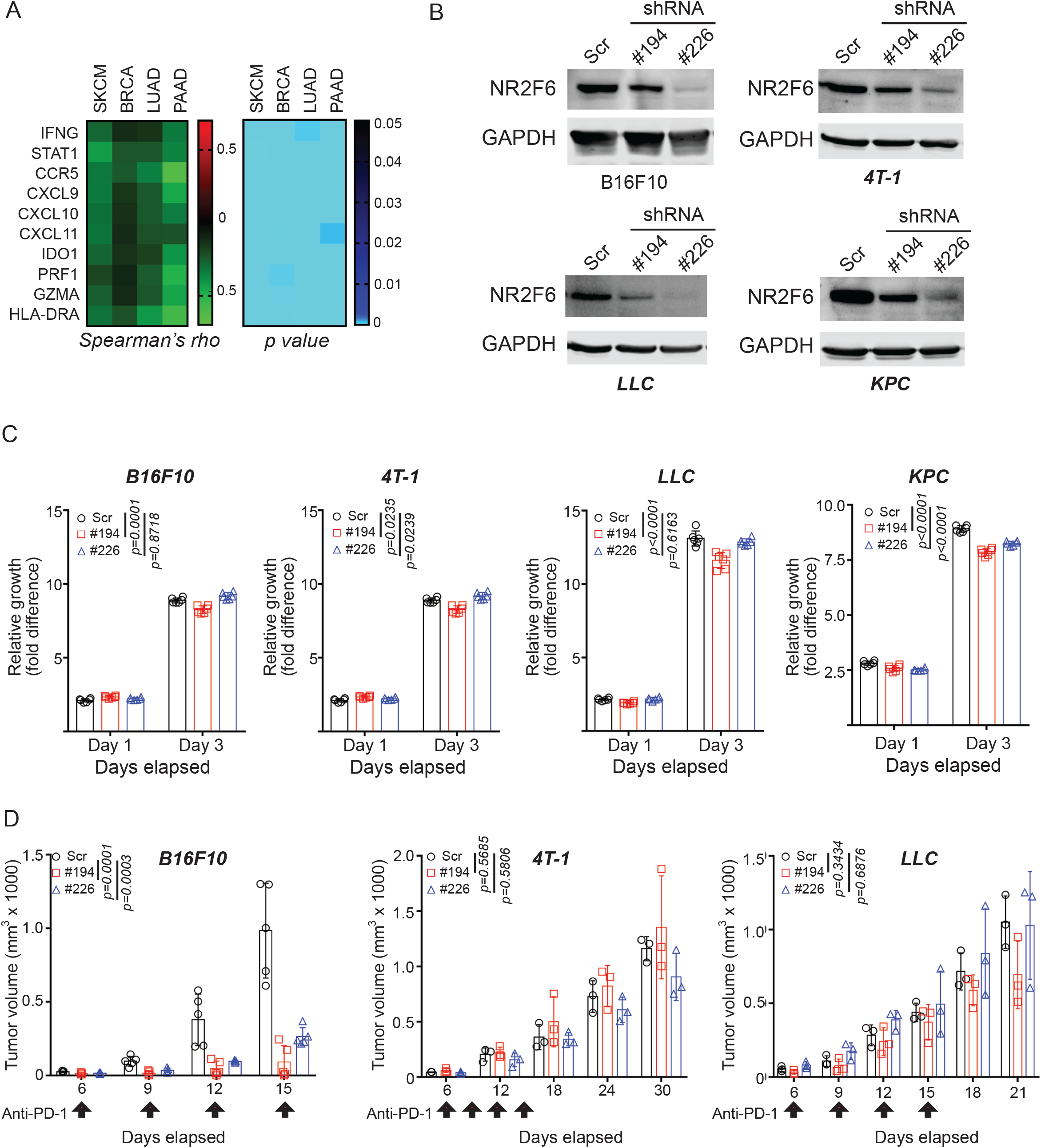
NR2F6 control of antitumor immunity in different cancer types. (A) Heatmaps show Spearman correlation coefficient (left) and corresponding p values (right) for NR2F6 expression (X-axis) in the indicated cancer types [melanoma (SKCM), breast (BRCA), lung (LUAD) and pancreatic (PAAD) cancers) with 10 IFNγ−signature genes (Y-axis). (B) Murine models for the cancer types [B16F10 (SKCM), 4T-1 (BRCA), LLC (LUAD) and KPC (PAAD)] in (A) were transduced with scrambled (Scr) or two shRNAs (#194 and #226) against NR2F6. The KD efficiency was assessed by immunoblotting using indicated antibodies. (C) The growth of transduced cells *in vitro* was assessed by CellTiter-Glo. Relative fold-differences in luminescence at days 1 and 3 were calculated relative to luminescence on day 0, defined arbitrarily as 1. (D) The transduced cells were inoculated to C57BL/6 (B16F10 and LLC) or BALB/C (4T-1) mice, which were treated with anti-PD-1 antibody (RMP1-14) at days 6, 9, 12, and 15 (arrows). Tumor volumes were monitored at indicated time points.

**Supplementary figure S4.**
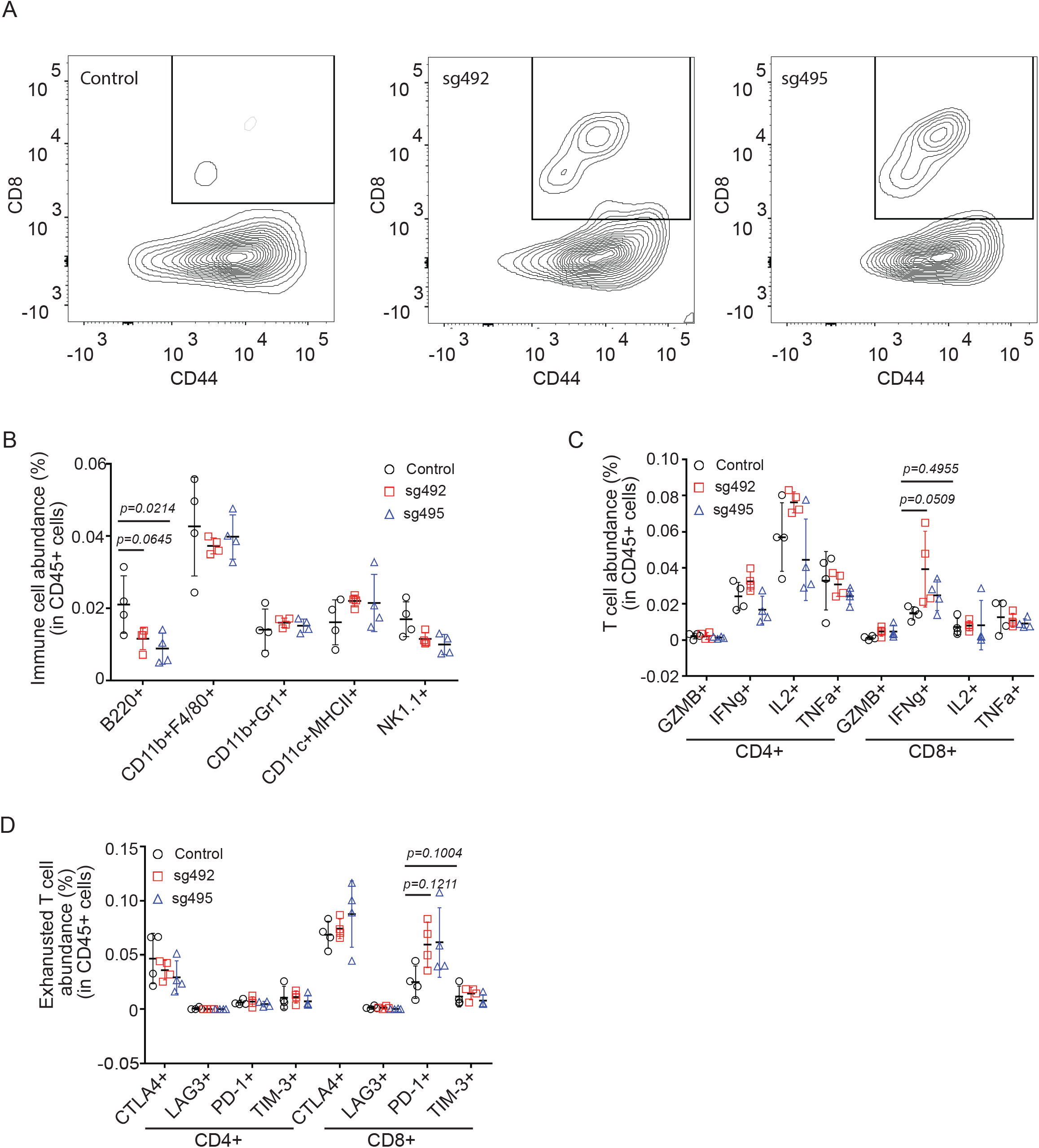
Immune cell abundance in NR2F6 KO tumors. (A) Representative contour plots in Figure 3C. (B) The abundance of immune cell types in CD45^+^ cells was assessed by FACS using indicated surface markers [B cells (B220^+^), macrophage (CD11B^+^F4/80^+^), MDSC (CD11b^+^Gr1^+^), dendritic cells (CD11c^+^MHCII^+^), and NK (NK1.1^+^)]. (C) The abundance of T cells producing indicated cytokines in CD45^+^ cells was assessed by FACS. (D) The abundance of exhausted T cells in CD45^+^ cells was assessed by FACS.

**Supplementary figure S5.**
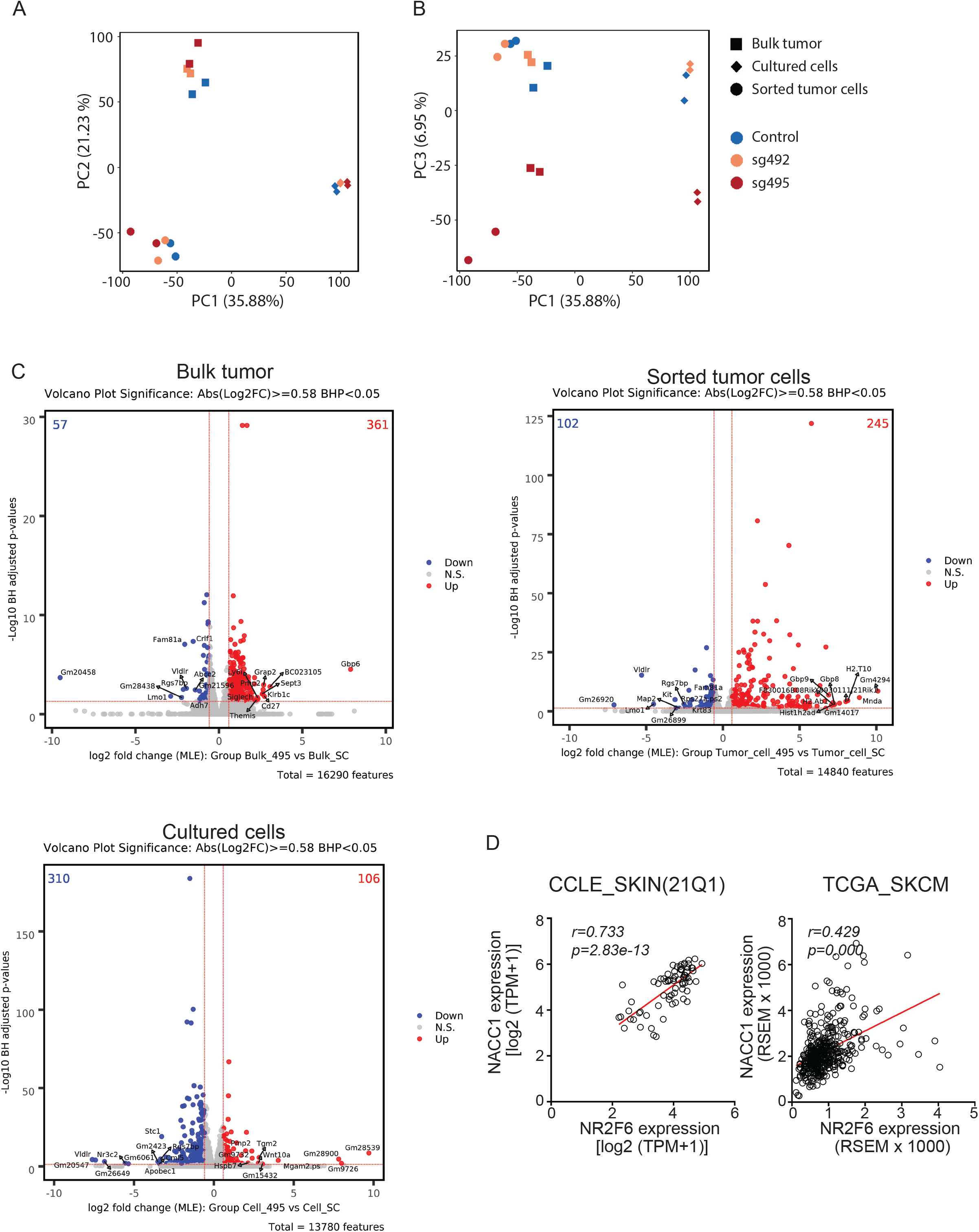
Identification of putative NR2F6 effectors. (A-B) RNAseq was carried out as described in Figure 4. Principal component analysis (PCA) of RNAseq data was visualized. (C) Volcano plots visualize up- or down-regulated genes in RNAseq analyses of bulk, MACS-sorted, and cultured cells. (D) The expressions of NR2F6 and NACC1 in human melanoma cell lines (Emran *et al*.) and human melanoma tissues (TCGA) were assessed. Pearson correlation coefficient (*r*) and corresponding p values (*p*)were presented.

**Supplementary figure S6.**
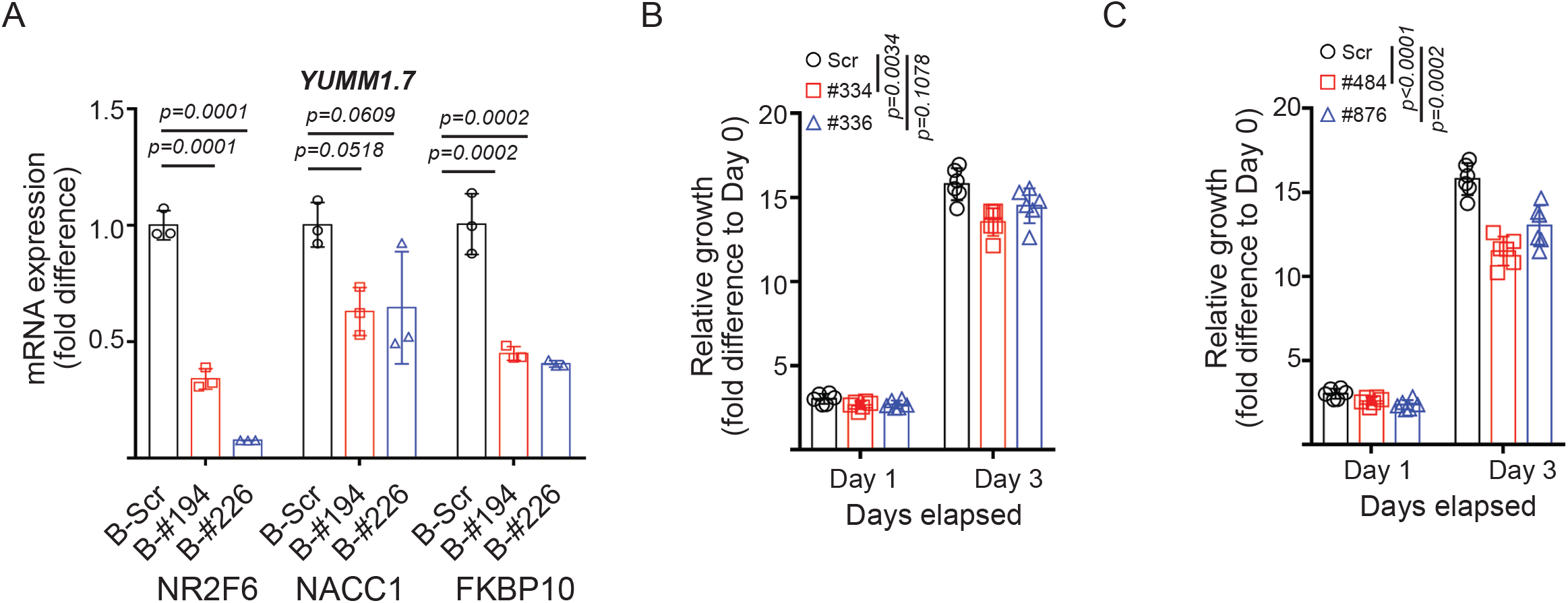
NACC1 and FKBP10 as downstream NR2F6 effectors. (A) The expressions of indicated genes were assessed in YUMM1.7 (B-C) Growth of transduced cells with indicated shRNAs, shNACC1 (B) and shFKBP10 (C), were assessed using CellTiter-Glo. Relative fold-differences in luminescence at days 1 and 3 were calculated relative to luminescence on day 0, defined arbitrarily as 1.

